# Neuroimmune mechanisms of a mouse model of chronic back pain

**DOI:** 10.1101/2024.12.13.628454

**Authors:** Aleyah E. Goins, Nesia A. Zurek, Cristian O. Holguin, Alexis Gravelle, Sachin Goyal, Shahani Noor, Jenna B. Demeter, Maddy R. Koch, June Bryan I. de la Peña, Karin N. Westlund, Sascha R.A. Alles

**Affiliations:** Department of Anesthesiology and Critical Care Medicine, University of New Mexico Health Sciences Center, Albuquerque, NM, USA; Department of Neurosciences, University of New Mexico Health Sciences Center, Albuquerque, NM, USA

**Keywords:** Chronic back pain, Neuroimmune Mechanisms, Dorsal Root Ganglia Tissue, Sensory neurons

## Abstract

Chronic back pain (CBP) is the leading cause of disability affecting 1 in 10 people worldwide. Symptoms are marked by persistent lower back pain, reduced mobility, and heightened cold sensitivity. Here, we utilize a mouse model of CBP induced by injecting urokinase-type plasminogen activator (uPA), a proinflammatory agent in the fibrinolytic pathway, between the L2/L3 lumbar vertebrae. We identified neuroimmune interactions contributing to uPA-induced CBP (henceforth, uPA-CBP) in mouse dorsal root ganglia (DRG), where nociceptive neurons reside. Flow cytometric data reveal that uPA-CBP increases CD45+CD11b+ cells in the DRG, a population characteristically implicated in other chronic pain models^1^. Blocking colony stimulating factor 1 receptor (CSF1R) signaling using PLX5622 partially reduced pain, suggesting CD45+CD11b+ macrophage involvement. Whole-cell patch-clamp electrophysiology data indicated DRG neuron hyperexcitability in CBP mice compared to controls. RNA sequencing revealed upregulation of pain- and inflammation-related genes involved in leukocyte migration. Together, these findings underscore the importance of the DRG neuroimmune axis in mediating chronic back pain.

**Highlights:**

- uPA-CBP induces gait changes, mechanical and thermal sensitivity compared to shams
- uPA-CBP mice show increased CD45+CD11b+ cells in DRG compared to shams
- uPA-CBP mice show neuronal excitability in DRG neurons compared to shams
- Pain behaviors are alleviated by pharmacologically blocking CSF1R signaling
- Dysregulation of inflammation- and ion channel-related genes in uPA-CBP DRG

## Introduction

According to the NIH Pain Consortium^2,3^, chronic pain affects 1 in 5 people worldwide and about half of those cases are chronic back pain (CBP). CBP commonly limits activity by causing persistent hypersensitivity to touch, temperature, and pressure. It has been discovered that these symptoms are driven in part by observable physiological changes such as neuroinflammation, maladaptive tissue remodeling, and sensitization of nociceptive pathways in the peripheral nervous system (PNS)^4^. The dorsal root ganglia (DRG) house the cell bodies of peripheral sensory neurons responsible for detecting and relaying environmental stimuli to the central nervous system (CNS). Damage to the DRG, or along its distal axon, can disrupt these signals, leading to numbness, pain disorders, or other altered sensations^5^. Importantly, DRGs are subject to peripheral immune cells. In particular, macrophages proliferate and differentiate into pro-or anti-inflammatory phenotypes in the DRG, especially in pain states^6^. Several inflammatory mediators such as chemokine (CC motif) ligand 2, interleukin (IL)-6, IL-1β, matrix metallopeptidases-2 (MMP2), and -9 (MMP9), and CX3CL1 are indicators of nerve injury^7^ and propagate nociception through macrophage activation^4,8^.

Here, we describe a novel uPA-CBP mouse model induced by a single injection of urokinase-type plasminogen activator (uPA) into the L2-L3 spinous ligament. uPA has been established in our previous studies as a sensitizer^9,10^ but peripheral neuroimmune contributions to pain were unknown. Behavioral assessments revealed that male and female uPA-CBP mice displayed significant mechanical hypersensitivity, cold hypersensitivity, and gait disturbances, indicating increased pain behaviors compared to sham controls. Electrophysiological analysis of DRG neurons in male uPA-CBP mice demonstrated increased neuronal excitability, characterized by altered input resistance, increased firing frequency, and heightened depolarizing spontaneous fluctuations (DSFs) in prevalence, frequency, and amplitude. At the immune level, altered monocyte production and maturation significantly influenced the development of pain behaviors and neuronal excitability in uPA-CBP mice, persisting up to eight weeks post-model induction. Flow cytometry and immunohistochemistry confirmed macrophage involvement, with blocking colony-stimulating factor 1 receptor (CSF1R) signaling via PLX5622 alleviating pain behaviors, implicating CD45+CD11b+ macrophages in CBP pathophysiology. Transcriptomic profiling of lumbar DRGs identified 18 differentially expressed genes (DEGs) and over-represented canonical pain- and immune-related enriched pathways in uPA-CBP compared to sham controls. These findings suggest novel therapeutic targets within the DRG neuroimmune axis, offering potential interventions that target peripheral immune cell populations to treat CBP.

In summary, this study provides a comprehensive characterization of the uPA-CBP model and highlights the importance of neuroimmune interactions in CBP, paving the way for novel therapeutic strategies. While we acknowledge that clinical and physiological presentation of pain and immune markers can be highly variable between sexes^11^, we have focused on male mice for our studies but identifying sex differences will be a subject of future study.

## Methods

### Urokinase plasminogen type activator-induced chronic back pain model (uPA-CBP)

All animal procedures described followed the NIH Guide for the Care and Use of Laboratory Animals. Studies were approved by the local Institutional Care and Use Committee (IACUC #23-201364-HSC) of the University of New Mexico Health Sciences Center. All studies complied with policies under the auspices of an OLAW Assurance of Compliance (A3002-01) on the use of animals in research, as described in Part III. II. Assurances and Certifications. Animals were housed in the Animal Resources Center (ARC) housing facility maintained by the laboratory staff and Division of Laboratory and Animal Resources (DLAR) staff. uPA-CBP model induction methods were previously published in Montera et al., 2023^9^. Male and female BALB/c mice (Envigo Labs) were 8 to 10 weeks old at start of experiments. All animals were injected in the L2-L3 spinous ligament region via Hamilton syringe with 5 microliters (µL) of either vehicle (0.9% saline) or urokinase-type plasminogen activator (uPA, 2mg/ml, CC4000, Millipore Sigma) under anesthesia (4% isoflurane and 2 liters of oxygen). Following injections, mice were placed in a warmed recovery chamber until full mobility returned (<30 seconds). Mouse weight was checked weekly and underwent *in vivo* behavioral testing. Then the mice were sacrificed for experiments 8 weeks post-model induction.

PLX5622 (MedChemExpress, Cat# HY-11415) was used to pharmacologically inhibit CSF1R signaling. Recipe was determined from Riquier and Sollars, 2020^12^. PLX5622 was made at a concentration of 50mg/kg in a vehicle containing 5% DMSO, 20% Kolliphor RH40, and 0.01M phosphate buffer solution (PBS, Cat #14-200-166, ThermoFisher). For 3 days, mice received i.p. injections twice per day (12 hours apart) with either PLX5622 or vehicle alone. Then, the mice underwent uPA-CBP model induction after 5 days.

### Behavioral assays

Behavioral assays were performed as described in Montera et al., 2023^9^. Male and female BALB/c mice were tested at baseline and subsequently every two weeks for 8 weeks for mechanical sensitivity. The mice were tested at 4-weeks and 8-weeks for cold and heat sensitivity, also at 8-weeks for gait disturbances. Mechanical sensitivity was quantified using von Frey filaments (Bioseb) with the up-down method to determine the threshold at which a stimulus elicits a paw withdraw response. Cold and heat sensitivity were assessed on a cold/hot plate device (IITC Life Science Incremental Hot/Cold Plate) by measuring paw withdraw latency in seconds (s). Withdraw latency was defined as the time from applying stimulus to the mouse eliciting a positive response indicated by biting, flicking, or repeatedly lifting the inflicted hind paw. Cold plate was set to gradually decrease temperature until -9°C. Tests were terminated when a positive response was elicited or the temperature reached -9°C. Two trials were taken for each mouse given by two different experimenters with 1 minute rest period in between trials. The Hargreaves test applies a constant source of heat to a hind paw with an infrared laser. Tests was terminated after 15 seconds or when a positive response was elicited. Five trials were taken for each mouse with 1 minute rest period between trials. Gait disturbances (significant differences in gait length and width) were measured with ink blots of hind paw prints and were analyzed in FIJI (NIH).

### DRG tissue preparation

Mice were anesthetized (4% isoflurane and 2 liters of oxygen) until there was no reaction to a toe pinch by forceps. Six DRGs were extracted from mouse lumbar enlargement (levels L2-L4) via laminectomy. Each DRG was gently removed with fine-tip forceps from the vertebral column. DRGs were collected in Mg^2+^- and-Ca^2+^-free Hank’s Buffered Saline Solution (HBSS) 1x buffer. The tissue was transferred to RNA-later for RNA sequencing. For flow cytometry and whole-cell patch-clamp electrophysiology experiments, the tissue was enzymatically dissociated with dispase II and collagenase type 2 to separate cells from connective tissues for 30 minutes at 37°C in an incubator with 5% CO_2_. Gentle agitation was done every 10 minutes with a pipette tip. Further mechanical dissociation was done using trituration with a flame polished glass pipette to ensure sufficient cell separation. When no pieces of tissue were visible in the media, the cell suspension was filtered with a 100-micron cell strainer with Dulbecco’s Modified Eagle Medium (DMEM) supplemented with 10% fetal bovine serum and 1% antibiotic-antimycotic (CatCat #15240062, GIBCO). The cell suspension was centrifuged with supplemented DMEM media. The cell pellet was resuspended in either Fluorescence Activated Cell Sorting (FACS) buffer (0.75% bovine serum albumin in PBS 1x, no phenol red, no salts) to be stained accordingly for flow cytometry or in supplemented DMEM to be plated on a glass coverslip for primary cell culture.

### Flow cytometry

The dissociated cell suspension in FACS buffer was prepared for fluorescent staining with antibodies against cell surface and intracellular markers. Equal cell counts between samples was established. Unstained control and viability positive control tubes were prepared from sample tubes. All fluorochrome conjugated antibodies were purchased from ThermoFisher (**Fig. 2A**). Fc block (1:10, Cat #553142, BD Bioscience) was applied for 10 minutes in all tubes prior to adding antibodies. The unstained control tube underwent all steps without receiving fluorochrome conjugated antibodies or viability dyes. The viability positive control tube was boiled at 65°C for 5 minutes to induce cell death and then Fixable Viability Dye was added for 1 hour. The sample tubes received fluorochrome conjugated antibodies against cell surface markers (CD45 (Cat #48-0451-82), CD3 (Cat #12-0031-82), CD11b (Cat #64-0112-82), CD14 (Cat #46-0141-82)) at the dilutions listed in **Figure 2A**. All tubes were washed with FACS buffer and then prepared for fixation and permeabilization with an intracellular staining kit (Cat #88-8824-00, ThermoFisher). The sample tubes received fluorochrome conjugated antibodies against intracellular markers (GFAP (Cat #53-9892-82), TNFα (Cat #17-7321-82)) at the dilutions listed in **Figure 2A**. Simultaneously, tubes for compensation beads were prepared with a single color each. Then all tubes underwent fluorescent detection with an Attune NxT Acoustic Focusing Cytometer (ThermoFisher). The Flow Cytometry Standard (.fcs) files were analyzed in FlowJo™ software (v10.10, Becton, Dickinson and Company, OR). Events were restricted by randomized algorithm to normalize cell count across samples. Gating strategy was designed to exclude debris, doublets, and dead cells (**Fig. 2B**). Negative and positive populations were distinguished with the use of spectral-enabled compensation calculations automated by FlowJo™ for multi-color detection. After gating on the intended cell population, the target marker’s percentage of the parent population is taken to calculate total cell count of the target marker and the positive percentage of all live singlets in Microsoft Excel.

### Whole-cell patch-clamp electrophysiology

Primary mouse DRG neuronal cultures were used to observe neuronal excitability with whole-cell patch-clamp electrophysiology in sham and uPA-CBP mice. Recordings were conducted 1 day *in vitro* (DIV1) at room temperature in an artificial cerebrospinal fluid (aCSF)-perfused chamber, recipe found in previous publications^13^. Neurons were identified using differential interference contrast optics connected to an IR-2000 digital camera (Dage MTI, IN). Cells with low resting membrane potential (above -35 millivolts (mV), not corrected for liquid junction potential) or with high series resistance (>15 megaohms (MΩ)) were omitted from analysis. Nociceptors were identified by size of neurons being small to medium in diameter (20-30 microns)^14^. Current and voltage clamp recordings utilized a Molecular Devices Multiclamp 700B amplifier, with signals filtered at 5 kHz, acquired at 50 kHz, and recorded using Clampex 11.2 software. Depolarizing spontaneous fluctuations (DSFs) were compared between conditions following quantitative analysis with the automated FIBSI program which detected frequency and amplitude of DSF peaks for each neuron^15^. Methods for frequency over current (f-I) plot generation from Zurek et al., (2024) were used^16^. Analysis was performed in Easy Electrophysiology v.2.5.1, Clampfit 11.2 (Molecular Devices), JupyterLab, matplotlib, PANDAS and the Python v3.12 package pyABF. Code for analysis and package citations can be found here: https://github.com/AllesLab/Goins-et-al-Neuroimmune-2024.git

### Immunohistochemistry

Methods were modified from Abcam’s “complete IHC guide”^17^. Eight weeks post-model induction, sham and uPA-CBP were anesthetized (4% isoflurane, 2 liters of oxygen) and transcardial perfusion fixed with 4% paraformaldehyde (PFA) (Cat #50-980-494, Fisher Scientific) in PBS 1x until stiff. DRGs were extracted and placed in 4% PFA overnight at 4°C and moved to 30% sucrose until tissue sank. Then, the DRGs were embedded in OCT (Cat #25608-930, VWR), sectioned on a cryostat at 16 microns thick, and allowed to adhere to glass slides overnight. On a platform rocker, the slides were gently washed with PBS and permeabilized with PBST (PBS + 0.025% Triton-X-100 (Cat #X100-100ML, Millipore Sigma)). The slides were blocked in PBS + 10% bovine serum albumin (BSA) (Cat #10735078001, Millipore Sigma). Diluted primary antibodies or fluorophore conjugated antibodies from **Figure 2A** were applied to the slides with 1% BSA in PBS for overnight incubation at 4°C. Then, the slides were washed in PBS and further permeabilized in PBST. On a platform rocker, diluted secondary antibodies were applied with 1% BSA in PBS to the slides in a dark humidity chamber for 2 hours at room temperature. After washing with PBS, the slides were mounted with Fluoromount-G with DAPI (Cat #00-4959-52, ThermoFisher). Confocal imaging was done on an Olympus IXplore Spin/Yokogawa CSU-W1 Confocal Microscope at 10X and 60X (Evident Microscopy, Waltham, MA). Two staining panels were selected to demonstrate distinct cell populations in DRG tissue. The first panel (scale bar=100 microns) was stained for peripherin (1:500, Cat #PA1-10012, Invitrogen) visualized with secondary antibody AF555 (1:2000, Cat # AB150170, Abcam), F4/80 conjugated with AF488 (1:100, Cat #MF48020, ThermoFisher), and CD11b conjugated with SB645 (1:100, Cat #64-0112-82, ThermoFisher). The second panel (scale bar=50 microns) was stained for peripherin (1:500) visualized with secondary antibody AF555 (1:2000), GFAP conjugated with AF488 (1:250, Cat #53-9892-82, ThermoFisher), and CD45 conjugated with PE-Cy7 (1:100, Cat #48-0451-82, ThermoFisher).

### RNA sequencing and bioinformatic analysis

Six lumbar DRGs each were extracted from sham and uPA-CBP mice and stored in RNAlater. Total RNA was isolated using the Monarch Total RNA MiniPrep Kit (Cat #T2010S, New England Biolabs) following the manufacturer’s protocol. RNA quality was ensured using a NanoDrop spectrophometer (ThermoFisher) and a 5400 Fragment Analyzer System (Agilent). During library preparation, oligo(dT) beads were used to enrich mRNAs. Libraries were sequenced on a NovaSeq X Plus platform (Illumina, 150 bp paired-end reads, >20 million read pairs/sample) (Novogene). Phred scores (Q20 > 97.8% and Q30 > 93.8% for all samples) indicated high read quality. Paired reads were filtered out if either read 1) was contaminated with an adapter, 2) had > 10% uncertain nucleotides (N) (rarely), or 3) was of low quality (Q5 for > 50% of nucleotides). Read quality was further ensured using FastQC. As an artifact of two-color chemistry systems, Poly-G tails ≥20 bp were trimmed using BBDuk. HISAT2 was used to align the reads to the Ensembl reference genome (GRCm39, release 112), and StringTie was used to quantify gene expression. Differential expression analysis was conducted in edgeR, in which raw gene counts were normalized using the trimmed mean of M-values (TMM) method, and the quantile-adjusted conditional maximum likelihood (qCML) method was used to assess differential expression of genes. Differentially expressed genes (DEGs) were identified as those with P value ≤0.05, |log2FC| ≥ 0.5, and false discovery rate (FDR) ≤0.05 calculated with Benjamini-Hochberg corrections method. Genes with |log2FC| ≥0.5 and a p-value ≤0.05 were further analyzed for biological significance through an over representation analysis with Gene Ontology (GO) terms, specifically biological processes, cellular components, and molecular functions.

We used a calculated GO enrichment percentage to measure the number of genes associated with each GO term in both the input (test) gene set and the background gene set. The input gene set included the genes of interest, while the background set represents all genes considered in the analysis. For each GO term, we calculated two ratios: The proportion of genes in the input set associated with the GO term (GeneRatio) and the proportion of genes in the background set associated with the same GO term (BgRatio). The enrichment ratio was then calculated by dividing the GeneRatio by the BgRatio. This value shows how prevalent a GO term is in the test gene set compared to the background genes. To express this enrichment as a percentage, the ratio was multiplied by 100. The R packages used included edgeR, limma, statmod, clusterProfiler, org.Mm.eg.db, AnnotationDbi, GO.db, topGO, enrichplot, tinyverse, and ggplot2. Code for analysis and package citations can be found here: https://github.com/AllesLab/Goins-et-al-Neuroimmune-2024.git

### Statistical significance tests and data visualization

A p-value of <0.05 was considered statistically significant and tests used are indicated in the figure legends and text below. Analysis was performed using GraphPad Prism v10.0.2 unless otherwise stated above. Error bars denote mean ± standard error of the mean (SEM).

## Results

### Persistent hypersensitivity and gait disturbances characterize uPA-CBP model

Male (n=13 per group) and female (n=4 per group) BALB/c mice were placed in either sham or uPA-CBP groups and then underwent behavioral assays to assess presence of pain behaviors. A lower threshold to mechanical stimulus before eliciting a positive response indicates increased mechanical hypersensitivity; measured by von Frey application. Male uPA-CBP mice displayed persistent mechanical hypersensitivity over 8 weeks post-model induction compared to shams (column effect p<0.001, mixed-effects analysis). More specifically, 6 weeks-(p<0.01, Tukey’s multiple comparisons) and 8 weeks-(p<0.01, Tukey’s multiple comparisons) post-model induction, uPA-CBP exhibited a significantly lower threshold to mechanical stimulus compared to shams (**Fig. 1A**). Female uPA-CBP mice similarly showed persistent mechanical hypersensitivity over 8 weeks post-model induction compared to shams (column effect p<0.0001, mixed-effects analysis). More specifically, 4 weeks-(p<0.05, Tukey’s multiple comparisons), 5 weeks-(p<0.01, Tukey’s multiple comparisons), 7 weeks-(p<0.05, Tukey’s multiple comparisons), and 8 weeks-(p<0.05, Tukey’s multiple comparisons) post-model induction, female uPA-CBP mice presented a significantly lower threshold to mechanical stimulus compared to shams (**Fig. 1B**).

**Figure 1.**
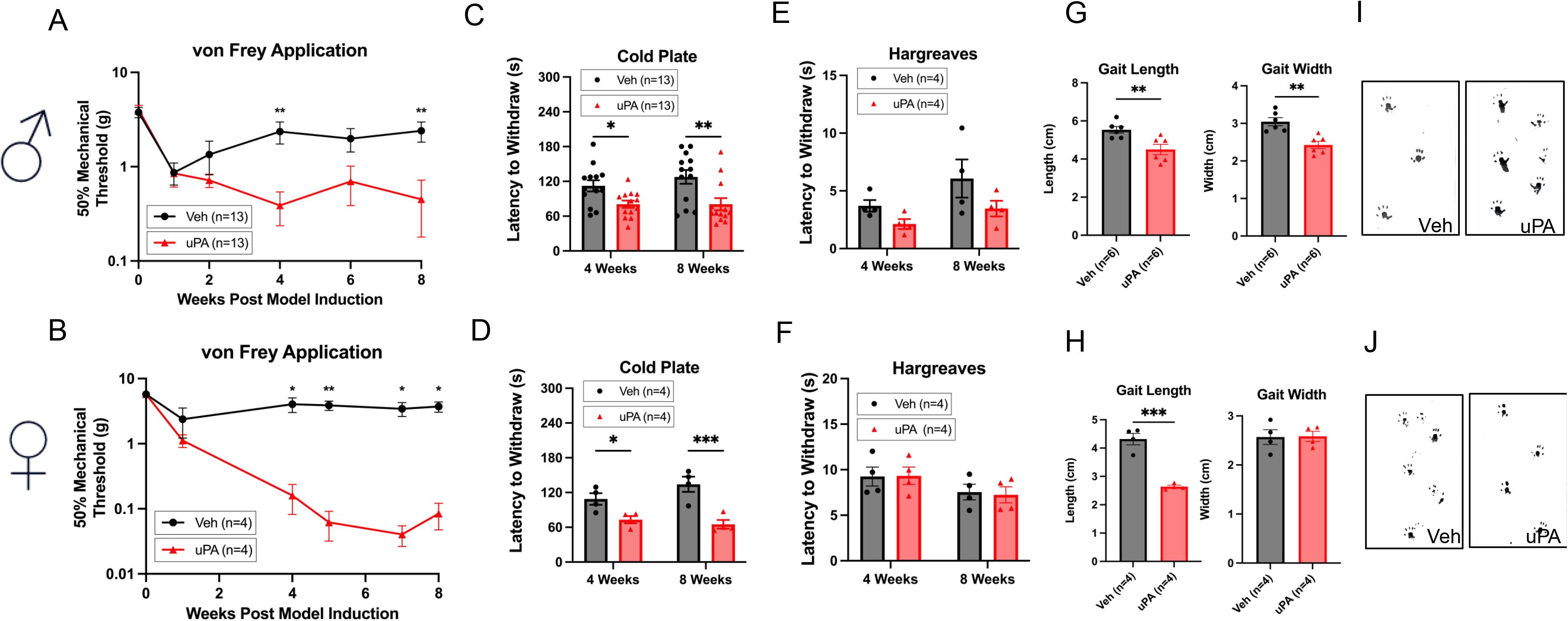
Behavioral assessment of pain-related responses and gait alterations in uPA-CBP and sham mice. Mechanical, cold, and gait-related pain behaviors were observed in uPA-CBP mice compared to sham controls. Mechanical hypersensitivity: (A) Male uPA-CBP mice (n=13) when compared to sham mice (n=13) exhibited persistent mechanical hypersensitivity over 8 weeks post-model induction compared to sham mice (column effect p<0.001, mixed-effects analysis). Specific differences were observed at 6 weeks (p<0.01, Tukey’s multiple comparisons) and 8 weeks (p<0.01, Tukey’s multiple comparisons). (B) Female uPA-CBP mice (n=4) when compared to sham mice (n=4) also displayed persistent mechanical hypersensitivity (column effect p<0.0001, mixed-effects analysis), with significantly lower thresholds at 4 weeks (p<0.05), 5 weeks (p<0.01), 7 weeks (p<0.05), and 8 weeks (p<0.05, Tukey’s multiple comparisons). Cold hypersensitivity: (C) Male uPA-CBP mice (n=13) when compared to sham mice (n=13) demonstrated significantly faster withdrawal responses to cold stimuli overall (column effect p<0.001, two-way ANOVA), with significant reductions in latency at 4 weeks (p<0.05, uncorrected Fisher’s LSD) and 8 weeks (p<0.001, uncorrected Fisher’s LSD). (D) Female uPA-CBP mice (n=4)when compared to sham mice (n=4) similarly had faster withdrawal responses overall (p<0.001, two-way ANOVA), with significant reductions in latency at 4 weeks (p<0.05) and 8 weeks (p<0.01, uncorrected Fisher’s LSD). Heat hypersensitivity: (E) Male uPA-CBP mice (n=4) when compared to sham mice (n=4) exhibited faster withdrawal responses to heat stimuli overall (p<0.05, two-way ANOVA), but there were no significant differences at 4 weeks (p>0.05, uncorrected Fisher’s LSD) or 8 weeks (p>0.05, uncorrected Fisher’s LSD). (F) Female uPA-CBP mice (n=4) when compared to sham mice (n=4) showed no significant differences in heat response latency overall (p>0.05, two-way ANOVA) or at either 4 weeks (p>0.05) or 8 weeks (p>0.05, uncorrected Fisher’s LSD). Gait analysis: (G) Male uPA-CBP mice (n=13) when compared to sham mice (n=13) had significantly shorter gait length (p<0.01, unpaired t-test) and width (p<0.01, unpaired t-test). (H) Female uPA-CBP mice (n=4) when compared to sham mice (n=4) had significantly shorter gait length (p<0.001, unpaired t-test) but no significant difference in width (p>0.05, unpaired t-test). (I, J) Representative ink blot images demonstrate gait alterations in male (I) and female (J) uPA-CBP mice compared to shams. Graph asterisks denote *p<0.05, **p<0.01, ***p<0.001, ***p<0.0001, ****p<0.00001.

Male (n=13 per group) and female (n=4 per group) BALB/c mice were assessed for cold hypersensitivity which was measured by latency to withdraw after being exposed to a cold stimulus at 4 weeks- and 8 weeks-post-model induction. Male uPA-CBP mice elicited faster positive responses than sham mice overall when the cold plate decreased in temperature (column effect p<0.001, two-way ANOVA). When examining each time point, male uPA-CBP mice had significantly lowered thresholds to cold stimulus at 4 weeks-(p<0.05, uncorrected Fisher’s LSD) and at 8 weeks-(p<0.001, uncorrected Fisher’s LSD) post-model induction (**Fig. 1C**). Female uPA-CBP mice also elicited generally faster positive responses than sham mice (column effect p<0.001, two-way ANOVA). At each time point, female uPA-CBP mice had significant reductions in latency to withdraw at 4 weeks-(p<0.05, uncorrected Fisher’s LSD) and at 8 weeks-(p<0.01, uncorrected Fisher’s LSD) post-model induction (**Fig. 1D**).

Male (n=4 per group) and female (n=4 per group) BALB/c mice were assessed for heat hypersensitivity which was measured by latency to withdraw after being exposed to a heat stimulus at 4 weeks- and 8 weeks-post-model induction. Male uPA-CBP mice elicited a faster positive response than sham mice overall when exposed to an infrared laser focused on a hind paw (column effect p<0.05, two-way ANOVA). However, male uPA-CBP mice did not have a difference in latency to withdraw at 4 weeks-(p>0.05, uncorrected Fisher’s LSD) or at 8 weeks-(p>0.05, uncorrected Fisher’s LSD) post-model induction (**Fig. 1E**). Female uPA-CBP mice did not elicit significantly different latency to withdraw than sham mice overall (column effect p>0.05, two-way ANOVA). Female uPA-CBP mice did not have significant differences in latency to withdraw at 4 weeks-(p>0.05, uncorrected Fisher’s LSD) or at 8 weeks-(p>0.05, uncorrected Fisher’s LSD) post-model induction (**Fig. 1F**).

Male (n=13 per group) and female (n=4 per group) uPA-CBP mice had their gait analyzed with ink blots of their hind paws across a paper sheet to evaluate stride patterns based on gait length and width. Male uPA-CBP demonstrated significantly shorter gait length (p<0.01, unpaired t test) and width (p<0.01, unpaired t test) when compared to sham mice (**Fig. 1G**). Female uPA-CBP mice showed significantly shorter length (p<0.001, unpaired t test) but not width (p>0.05, unpaired t test) when compared to sham mice (**Fig. 1H**). Representative images of the ink blots done by male mice shown in **Fig. 1I** and by female mice shown in **Fig. 1J**.

#### Monocyte-macrophage accumulate in DRG of uPA-CBP mice compared to sham mice

Flow cytometry and immunohistochemistry (IHC) were used in tandem to characterize non-neuronal cell populations within freshly isolated lumbar DRGs from male sham or uPA-CBP mice (n=4 per group, 6 DRGs per mouse). This approach enabled the simultaneous detection and analysis of neuronal, glial, and immune cell markers in DRG tissue with the use of fluorophore conjugated antibodies listed in **Figure 2A**. The gating strategy shown in **Figure 2B** was employed to identify viable singlet cells. Total cell count and percentage calculations were derived from the count and percentage of viable singlet cells of which were not significantly different between sham and uPA-CBP groups (p>0.05, unpaired t test) (**Fig. 2C**).

**Figure 2.**
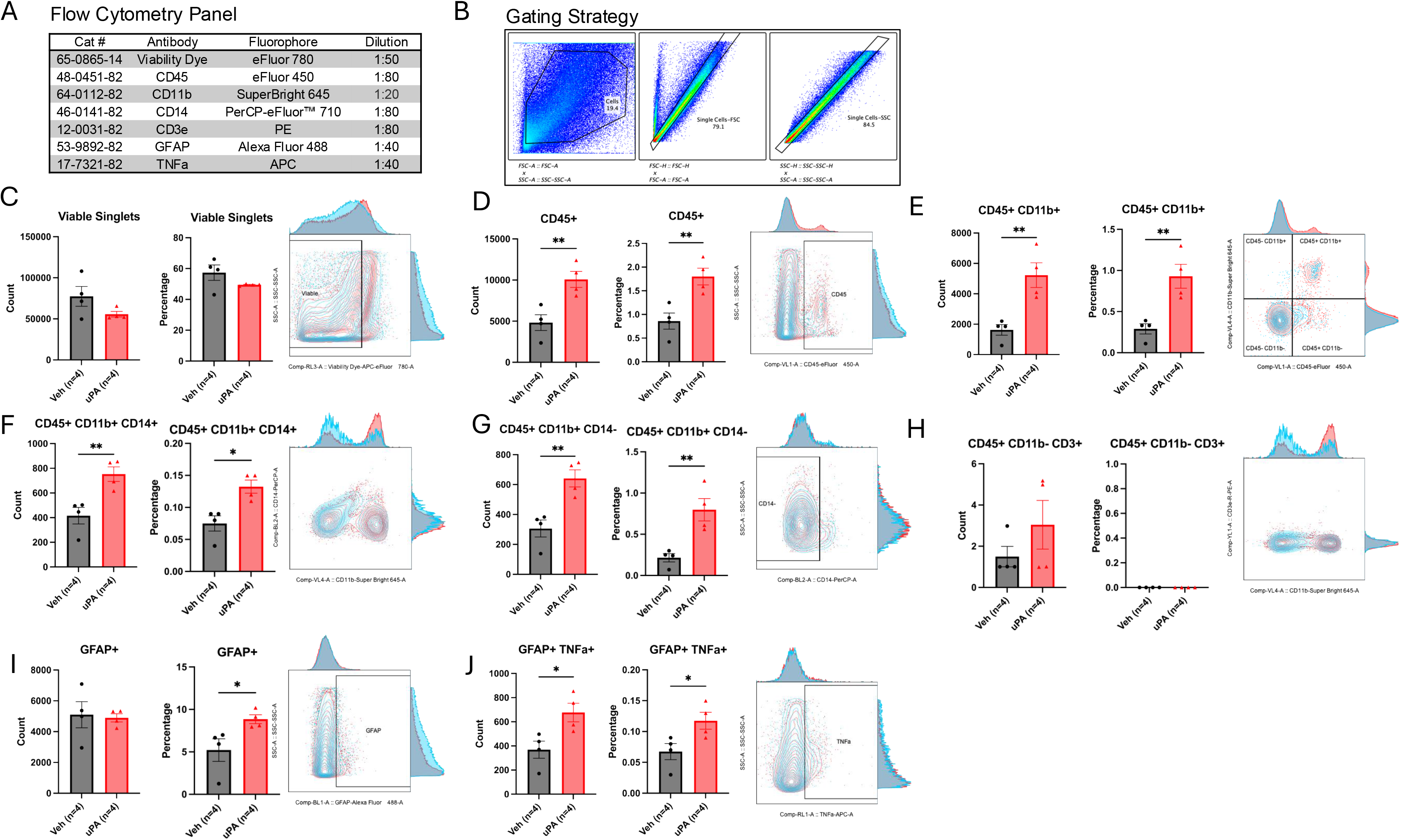
Flow cytometric characterization of non-neuronal cell populations in DRGs of sham mice (n=4) and uPA-CBP mice (n=4). (A) Fluorophore conjugated antibodies used for flow cytometric analysis of neuronal, glial, and immune cell markers in freshly isolated lumbar DRGs. (B) Gating strategy to identify viable singlet cells for further analysis of cell surface and intracellular markers. (C) Total cell count and percentage of viable singlet cells were not significantly different between sham and uPA-CBP mice (p>0.05, unpaired t-test). (D) CD45+ leukocyte count and percentage were significantly increased in the DRG of uPA-CBP mice compared to sham mice (p<0.01, unpaired t-test). (E) CD45+CD11b+ myeloid-monocyte cell count (p<0.01, unpaired t-test) and percentage (p<0.01, unpaired t-test) were significantly increased in the DRG of uPA-CBP mice compared to sham mice. (F) Monocyte-derived macrophages (CD45+CD11b+CD14+ cells) showed significantly increased count (p<0.01, unpaired t-test) and percentage (p<0.05, unpaired t-test) in uPA-CBP mice compared to sham mice. (G) CD45+CD11b+CD14-cells had significantly increased count (p<0.01, unpaired t-test) and percentage (p<0.01, unpaired t-test) in uPA-CBP mice compared to sham mice. (H) Lymphocytes (CD45+CD11b-CD3+ cells) were negligible and not statistically significant in count or percentage. (I) GFAP+ satellite glial cells (SGCs) had significantly increased percentage (p<0.05, unpaired t-test) but no significant differences in count (p>0.05, unpaired t-test). (J) Activated GFAP+TNFα+ SGCs showed significantly increased count (p<0.05, unpaired t-test) and percentage (p<0.05, unpaired t-test) in uPA-CBP mice compared to sham mice. Graph asterisks denote *p<0.05, **p<0.01, ***p<0.001, ***p<0.0001, ****p<0.00001.

Flow cytometric analysis revealed a significant increased count (p<0.01, unpaired t test) and percentage (p<0.01, unpaired t-test) of CD45+ leukocytes found in the DRG of uPA-CBP mice compared to sham mice (**Fig. 2D**). Furthermore, CD45+CD11b+ myeloid-monocyte cells were significantly increased in count (p<0.01, unpaired t test) and percentage (p<0.01, unpaired t test) in the DRG of uPA-CBP mice compared to sham mice (**Fig. 2E**). Moreover, monocyte-derived macrophages characterized by CD45+CD11b+CD14+ cells were significantly increased in count (p<0.01, unpaired t test) and percentage (p<0.05) in the DRG of uPA-CBP mice compared to sham mice (**Fig 2F**). In turn, neutrophils –albeit not exclusively– were identified by CD45+CD11b+CD14-were significantly increased in count (p<0.01, unpaired t test) and percentage (p<0.01, unpaired t test) in the DRG of uPA-CBP mice compared to sham mice (**Fig. 2G**). Presence of lymphocytes, detected by CD45+CD11b-CD3+ cells, was observed to be negligible and unfit for statistical significance in count and percentage in the DRG of uPA-CBP mice compared to sham mice (**Fig. 2H**). Satellite glial cells were specifically identified by GFAP and were observed to be significantly increased in percentage (p<0.05, unpaired t test), but had marginal cell count differences (p>0.05, unpaired t test) in the DRG of uPA-CBP mice compared to sham mice (**Fig. 2I**). Activated GFAP+ SGCs were detected with TNFα which were noted to be significantly increased in count (p<0.05, unpaired t test) and percentage (p<0.05, unpaired t test) in the DRG of uPA-CBP mice compared to sham mice (**Fig. 2J**).

IHC showcased abundant fluorescence of GFAP marker shown in cyan and CD45 marker shown in green by visual inspection of confocal imagery at 10x magnification in the DRG of uPA-CBP mice compared to sham mice (**Fig. 3A**). We observed GFAP expression in cells surrounding peripherin-labelled neurons (red) in the DRG of sham and uPA-CBP mice suggesting effectiveness for labelling SGCs (**Fig. 3A**). Additional markers were used to distinguish CD45+CD11b+ neutrophils from CD45+CD11b+ macrophages in a confocal image at 60x magnification (**Fig. 3B**). We found that CD11b expression colocalizes with F4/80 expression, a highly specific marker of macrophages (**Fig. 3B**). These results underscore significant non-neuronal cell involvement in the DRG of uPA-CBP mice compared to sham controls. In conclusion, we observed significant SGC activation, leukocyte accumulation in lumbar DRGs at 8 weeks post model induction of uPA-CBP compared to sham controls.

**Figure 3.**
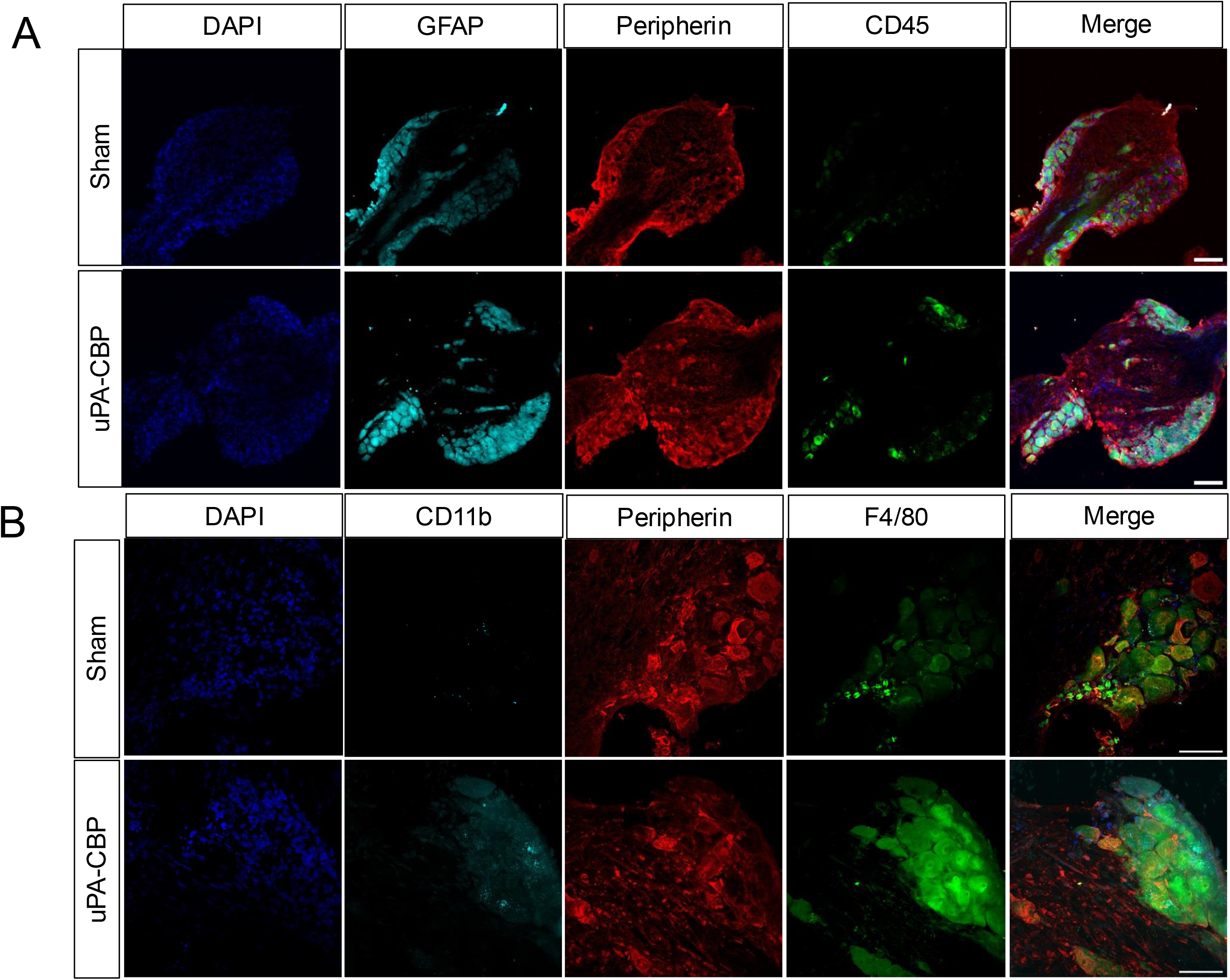
Immunohistochemical analysis of satellite glial cells (SGCs) and immune cell infiltration in DRGs from sham and uPA-CBP mice. (A) Confocal images at 10x magnification (scale: 100um) showing DAPI (blue), GFAP (cyan), peripherin (red), CD45 (green) markers in DRGs of sham and uPA-CBP mice. (B) Confocal images at 60x magnification (scale: 50um) showed DAPI (blue), CD11b (cyan), peripherin (red), F4/80 (green) markers in DRGs of sham and uPA-CBP mice. highlight significant non-neuronal cell involvement, including SGC expansion and macrophage accumulation, in the lumbar DRG of uPA-CBP mice compared to sham controls.

### Electrophysiological properties of DRG neurons are altered in uPA-CBP mice compared to sham mice

Whole-cell patch-clamp electrophysiology recordings were done in DRG neurons isolated from sham and uPA-CBP mice (n=4 per group) to determine the effects of the uPA-CBP model on DRG neuronal excitability and intrinsic properties. Rheobase measurements were not significantly different between sham (n=16 neurons) and uPA-CBP mice (n=19 neurons) (p>0.05, unpaired t test) (**Fig. 4A**). However, uPA-CBP mice (n=19 neurons) showed significant reduction in input resistance compared to sham mice (n=16 neurons) (p<0.05, unpaired t test) (**Fig. 4B**). The f-I relationship analysis showed that multi-firing DRG neurons from uPA-CBP mice (n=11 neurons) had significantly more APs at current injections over rheobase compared to sham mice (n=13 neurons) (column effect p<0.01, two-way ANOVA). Significant peak differences between uPA-CBP and sham neurons were observed at 60 pA (p<0.05, Šídák’s multiple comparisons), 70 pA (p<0.01, Šídák’s multiple comparisons), 80 pA (p<0.01, Šídák’s multiple comparisons), 90 pA (p<0.01, Šídák’s multiple comparisons), 100 pA (p<0.001, Šídák’s multiple comparisons), 110 pA (p<0.05, Šídák’s multiple comparisons, 120 pA (p<0.05, Šídák’s multiple comparisons), 140 pA (p<0.05, Šídák’s multiple comparisons), and 160 pA (p<0.05, Šídák’s multiple comparisons). The mean F-I slopes were 3.78 for uPA-CBP neurons and 2.38 for sham neurons (n=13), with a difference of -1.40 (SE=0.490, 95% CI: -2.42 to -0.384). The total area under the curve was larger for the slope of uPA-CBP neurons (mean ± SEM: 72.73 ± 6.632) compared to sham neurons (45.12 ± 4.443) (**Fig. 4C**). Representative f-I plots highlight the difference in AP firing rates between sham and uPA-CBP neurons, while also demonstrating presence of a rebound AP (**Fig. 4D**).

**Figure 4.**
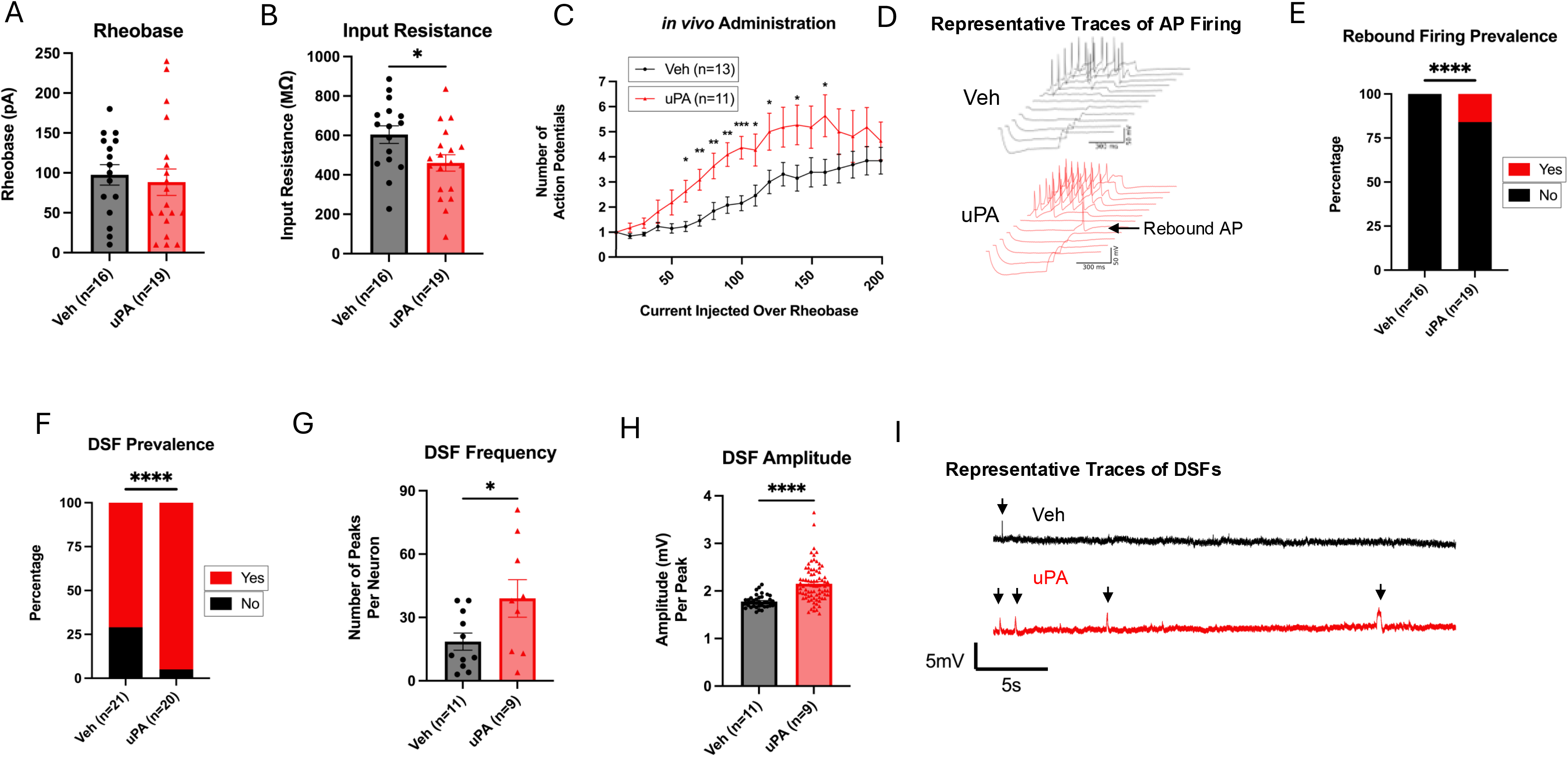
DRG neuronal excitability and intrinsic properties in sham and uPA-CBP mice. (A) Rheobase measurements were not significantly different between sham (n=16 neurons) and uPA-CBP mice (n=19 neurons) (p>0.05, unpaired t-test). (B) Input resistance was significantly reduced in uPA-CBP neurons compared to sham neurons (p<0.05, unpaired t-test). (C) Frequency-current (f-I) relationship analysis demonstrated significantly increased action potential (AP) firing in uPA-CBP neurons at current injections above rheobase (p<0.01, two-way ANOVA, column effect), with significant differences at 60 pA to 160 pA (p<0.05–0.001, Šídák’s multiple comparisons). The mean f-I slope and area under the curve were larger in uPA-CBP neurons. (D) Representative f-I plots show enhanced firing in uPA-CBP neurons and the presence of rebound APs. (E) Rebound prevalence was significantly higher in uPA-CBP neurons (16%) compared to sham neurons (0%) (p<0.0001, Fisher’s exact test). (F) DSF prevalence was significantly increased in uPA-CBP neurons compared to sham neurons (p<0.0001, Fisher’s exact test). (G) DSF frequency was significantly higher in uPA-CBP neurons (p<0.05, Mann-Whitney test). (H) DSF amplitude was significantly elevated in uPA-CBP neurons compared to sham neurons (p<0.0001, Mann-Whitney test). (I) Representative gap-free recordings illustrate DSF events in sham and uPA-CBP neurons. These findings collectively highlight increased excitability and altered intrinsic properties in uPA-CBP mouse DRG neurons. Graph asterisks denote *p<0.05, **p<0.01, ***p<0.001, ***p<0.0001, ****p<0.00001.

Rebound prevalence was significantly higher in uPA-CBP mice (n=19 neurons, 16% of cells, 3/19) compared to sham mice (n=16 neurons, 0% of cells, 0/16) (p<0.0001, Fisher’s exact test) (**Fig. 4E**). DSF prevalence was also significantly increased in uPA-CBP mice (n=20 neurons, 95%, 19/20) compared to sham mice (n=21 neurons, 71%, 15/21) (p<0.0001, Fisher’s exact test) (**Fig. 4F**). Depolarizing spontaneous fluctuations (DSF) frequency was significantly increased in uPA-CBP mice (n=9 neurons) compared to sham mice (n=11 neurons) (p<0.05, Mann-Whitney test) (**Fig. 4G**). DSF amplitude was also significantly elevated in the uPA-CBP mice (n=9 neurons) compared to sham mice (n=11 neurons) (p<0.0001, Mann-Whitney test) (**Fig. 4H**). Representative gap-free recordings that display DSFs are shown in **Figure 4I**. Overall, these findings underscore increased neuronal excitability in uPA-CBP mouse DRG neurons, evidenced by altered input resistance, increased firing frequency in f-I relationship, increased DSF prevalence, increased DSF frequency, and increased DSF amplitude.

### Characteristic pain behaviors and neuronal excitability of uPA-CBP are affected by CSF1R inhibition

To determine the role of macrophages in the development of uPA-CBP, male BALB/c mice were pretreated with PLX5622 (n=4 mice, hence described as “uPA-PLX5622”) or vehicle (n=5 mice, hence described as “uPA-veh”) before model induction. PLX5622 pretreatment significantly alleviated mechanical allodynia in uPA-CBP mice indicated by a higher mechanical threshold maintained up to 8 weeks post model induction (column effect p<0.0001, two-way ANOVA). The greatest reduction in sensitivity was seen at 8 weeks post-model induction in uPA-PLX5622 compared to uPA-veh mice (p<0.001, Tukey’s multiple comparisons) (**Fig. 5A**). However, uPA-PLX5622 mice were not significantly different from uPA-veh mice in sensitivity to cold or heat stimuli at 4 weeks (p>0.05, uncorrected Fisher’s LSD) or 8 weeks (p>0.05, uncorrected Fisher’s LSD) post-model induction (**Fig. 5B**). There were significant differences between uPA-veh and uPA-PLX5622 mice in gait length (p<0.001, unpaired t test), but not width (p>0.05, unpaired t test) (**Fig. 5C**). After behavioral testing, DRGs from uPA-veh mice (n=2 males) and uPA-PLX5622 mice (n=2 males) were used for whole-cell patch-clamp electrophysiology which revealed a significant reduction in DSF prevalence (p<0.001, Fisher’s exact test) in uPA-PLX5622 neurons (n=10, 30%, 3/10) compared to uPA-veh neurons (n=9, 56%, 5/9) (**Fig. 5D**). PLX5622 inhibits CSF1R which results in blocked production and maturation of monocytes, which are key to mononuclear phagocytic cell development such as macrophages^18,19^. We found that altered production and maturation of monocytes significantly affected the development of *in vivo* pain behaviors and *in vitro* DRG neuron excitability in uPA-CBP mice up to 8 weeks post model induction.

**Figure 5.**
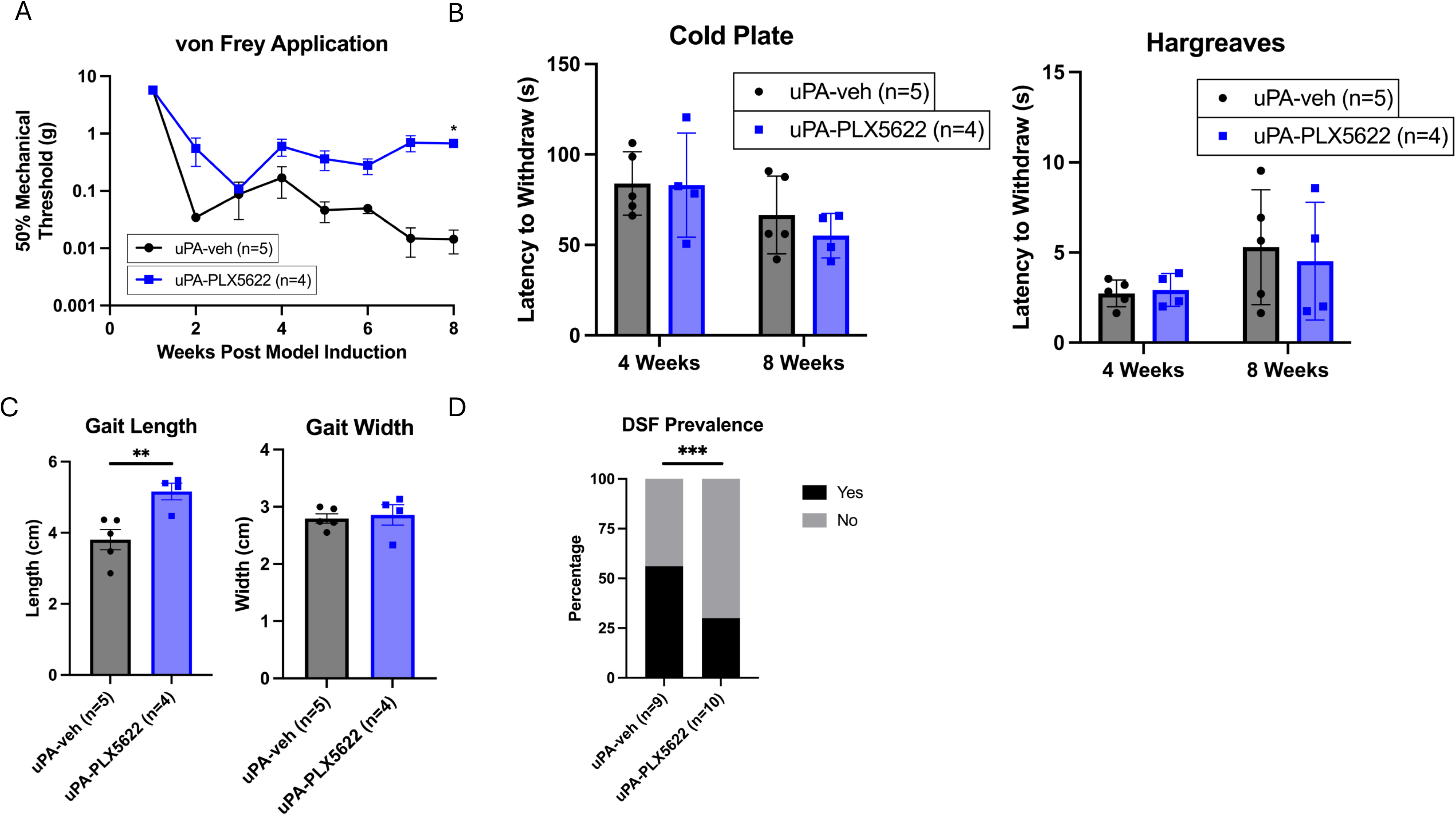
Effect of macrophage depletion via PLX5622 on pain behaviors and DRG neuronal properties in uPA-CBP mice. PLX5622-mediated inhibition of CSF1R disrupted monocyte maturation and macrophage development, reducing pain behaviors and neuronal excitability in the uPA-CBP model. (A) PLX5622 pretreatment (“uPA-PLX5622”, n=4) significantly alleviated mechanical allodynia compared to vehicle-treated uPA-CBP mice (“uPA-veh”, n=5), as indicated by a higher mechanical threshold maintained up to 8 weeks post-model induction (column effect p<0.0001, two-way ANOVA). The greatest reduction in sensitivity was observed at 8 weeks (p<0.001, Tukey’s multiple comparisons). (B) PLX5622 did not significantly affect sensitivity to cold (p>0.05, uncorrected Fisher’s LSD) or heat (p>0.05, uncorrected Fisher’s LSD) stimuli at 4- or 8-weeks post-model induction. (C) Gait length was significantly improved in uPA-PLX5622 mice compared to uPA-veh mice (p<0.001, unpaired t-test), but gait width was unaffected (p>0.05, unpaired t-test). (D) Whole-cell patch-clamp electrophysiology of DRG neurons revealed a significant reduction in DSF prevalence in uPA-PLX5622 neurons (30%, 3/10) compared to uPA-veh neurons (56%, 5/9) (p<0.001, Fisher’s exact test). Graph asterisks denote *p<.05, **p<0.01, ***p<0.001.

### DEGs related to immune and ion channel modulation are overrepresented in uPA-CBP mice compared to sham mice

Transcriptomic profiles were generated and statistically compared to identify DEGs in lumbar DRGs of uPA-CBP mice compared to sham mice (n=3 per group). RNAseq analysis elucidated the list of 18 significant DEGs, shown in ascending order of FDR (**Fig. 6A**). The significantly upregulated DEGs were *Arhgap15, Gm21986, Stk36, Klhl1, Gm21985, Ndor1, Gm38336, Rmi2, 2810454H06Rik, Mccc1os, Bco1, 4930544F09Rik, Ccdc63, Lrch4, Serpinb5*, and *Alg3*. The significantly downregulated DEGs were *Gm20186* and *Snx31*. There were 244 genes with p-values of ≤ 0.05 and |log2FC| ≥ 0.5 that were represented on the volcano plot but could not be determined as a significant DEG after benjamini-hochberg corrections (FDR≤0.05) in **Figure 6B**. Some notable genes consist of but are not limited to *Vgf, Nts, Trpa1, Gphn, Il1b, Pde4a, C3, Itgam, Csf3r, Lilrb4b, Il17d, Cd300a, Rftn1, Foxd1, Spi1, Gfap, Tlr9, Macf1, Mmp8, Adamts8, Col6a1, Stxbp5l, Macf1, Fbn1*, and *Plp1*. The 18 significant DEGs were labelled on the volcano plot with their respective gene name.

**Figure 6.**
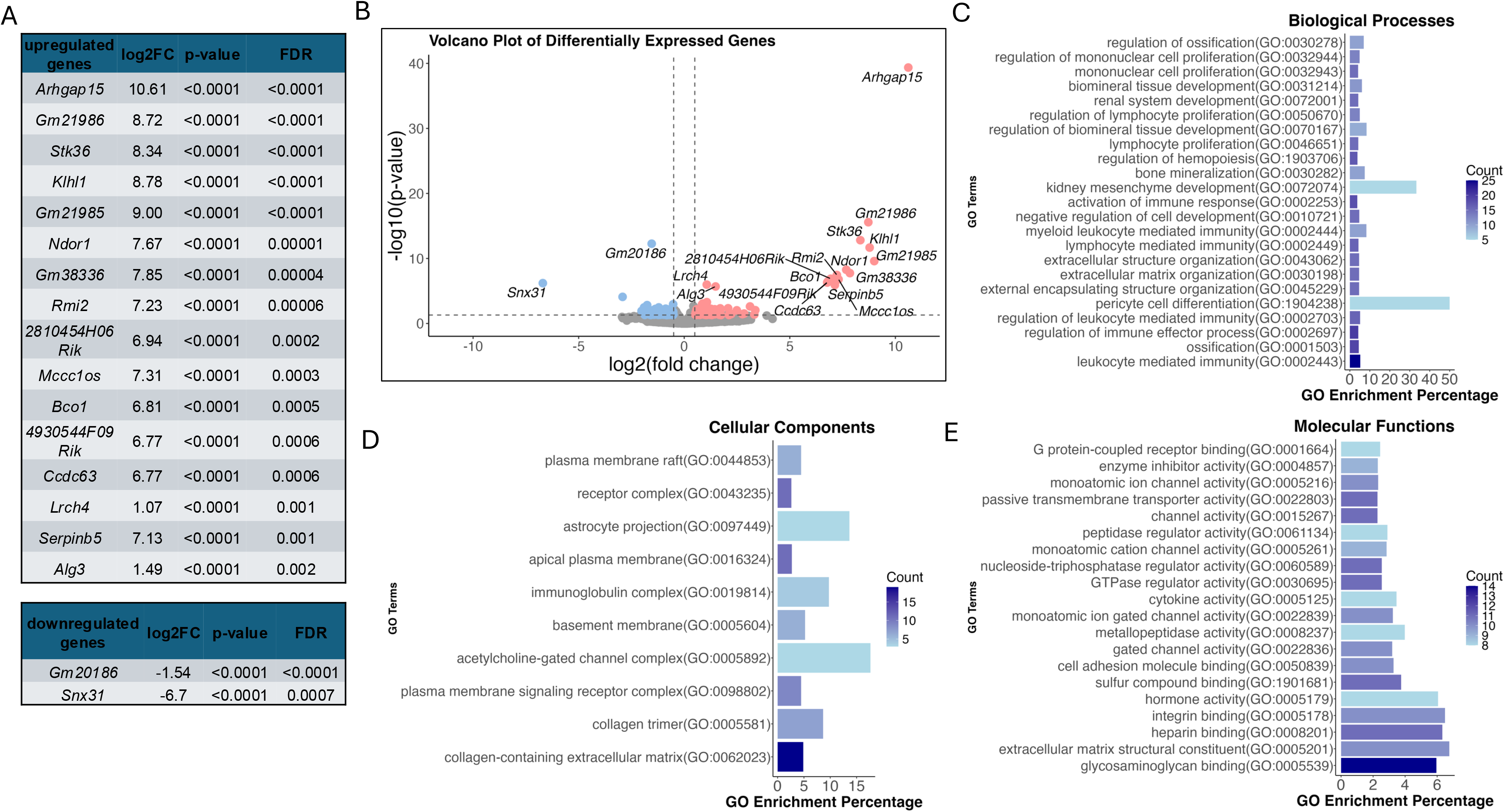
RNAseq reveals differential expression of pain- and inflammation-related genes and significant Gene Ontology (GO) enriched pathways in uPA-CBP model. Bulk RNAseq data obtained from DRG tissue extracted from sham and uPA-CBP male mice (n=3 per group). (A) Table of 18 significant differentially expressed genes (DEGs) produced by EdgeR analysis. Significance criteria were P-Value of ≤ 0.05, |log2FC| ≥ 0.5, and False Discovery Rate (FDR) ≤ 0.05. (B) Significance cut-offs were –log10(p-value)≤ 0.05 and log2FC≤-0.5 or ≥0.5; indicated by the grey dotted lines. Upregulated and downregulated genes are represented by red and blue points, respectively. Only genes with P-Value of ≤ 0.05, |log2FC| ≥ 0.5, and FDR < 1 are labelled. C-E) Top 20 significantly enriched GO terms from “biological processes”, “cellular components”, and “molecular functions” databases were sorted by q-value calculated from benjamini-Hochberg corrections and plotted in descending order. X-axis is the Go Enrichment Percentage. Color gradient represents the actual count of genes found in the GO term.

All 262 genes with p-values of ≤0.05 and |log2FC| ≥0.5 were considered for the GO enrichment pathway analysis. There were 896 enriched terms for biological processes, 56 for molecular functions, and 10 for cellular components. The top 20 enriched terms were plotted according to its percentage of all genes in the respective database and the color gradient indicates actual gene count in the pathway. GO terms leukocyte-mediated immunity (q<0.0001) and regulation of immune effector process (q<0.0001) had the greatest gene counts at 25 and 20 genes. Pericyte differentiation (q<0.0001) had the greatest GO enrichment percentage at 50% in the biological processes database (**Fig. 6C**). GO term collagen-containing extracellular matrix (q<0.014) showed the greatest gene count at 19 genes. Acetylcholine-gated channel complex possessed the greatest GO enrichment percentage at 17.6% in the cellular components database (**Fig. 6D**). GO term glycosaminoglycan binding (q<0.0001) had the largest gene count with 14 genes. Glycosaminoglycan binding (q<0.0001) had a GO enrichment percentage of 6% which marginally tied with extracellular matrix structural constituent at 6.8% (q<0.0001), integrin binding at 6.5% (q<0.0001), heparin binding at 6.3% (q<0.0001), and hormone activity at 6.1% (q=0.0006) of genes in the term category compared to the molecular functions database (**Fig. 6E**).

## Discussion

The primary findings of our experiments were that uPA, compared to vehicle injection, promoted altered pain behaviors and increased sensitivity which correlated with leukocyte recruitment to the DRG up to 8 weeks post-injection. We observed GFAP, CD45, CD11b, and F4/80 positive cell populations in the DRG tissue with immunohistochemistry. Furthermore, uPA-induced mechanical sensitivity was partially resolved by pharmaceutically inhibiting macrophage-colony stimulating factor 1 receptor (M-CSF1R) activity, indicating monocyte-derived cell involvement in the development of hypersensitivity. We performed whole-cell patch-clamp electrophysiology of DRG neurons from uPa-CBP mice and observed increased neuronal excitability including firing frequency and prevalence of DSFs compared to controls. We also performed bulk RNA-sequencing on the freshly extracted mouse DRG tissue and revealed enrichment of pathways associated with activated leukocyte mediated immune responses. In summary, we observed strong involvement of the DRG neuroimmune axis in our long-lasting model of CBP^1,20–22^.

### Pain- and immune-related characteristics of uPA-CBP mode

We demonstrated that the uPA-CBP model dysregulated a combination of genes associated with regulation of ion channel expression and immune system dynamics. For instance, *Klhl1* gene was upregulated in uPA-CBP mice compared to shams. Upregulated *Klhl1* contributes to Cav3.2 activity as well as rebound firing^23–25^. Our whole-cell patch-clamp electrophysiology findings suggest increased rebound action potentials which are regulated in part by T-type calcium currents, which are critical for neuronal excitability and pain perception^26^. *Klhl1* encodes the actin-binding protein KLHL1, which regulates T-type calcium currents by interacting with Cav3.2 channels, increasing current density and calcium influx^23^. T-type calcium channelosomes are promising targets of interest to address chronic pain disorders^27^. Irregular DSFs are produced by spontaneous background activity of diverse ion channels permeable to Na^+^ or Ca^2+^ combined with high somal input resistance^14^. They drive ongoing electrical discharge (>2mV peaks) measured in gap free current clamp recordings under painful conditions that increase nociceptor depolarization susceptibility^14^. Tian et al. (2024) found that frequency and amplitude of DSF-invoked peaks positively correlate with pain behaviors^14^. Our data were consistent with that assessment. We show that uPA-CBP mice have decreased input resistance and increased pain behaviors compared to shams. This may indicate that the uPA-CBP model does alter ion channel properties^14^, but more research is needed to determine the exact mechanism.

There were several upregulated genes that canonically participate in immune cell recruitment and activation. *Arhgap15* encodes a Rho GTPase-activating protein that promotes leukocyte migration^28^ and regulates neutrophil dynamics during inflammation^29^. *Bco1* encodes the enzyme that cleaves beta-carotene into vitamin A (retinal), a precursor for retinoic acid which regulates immune cell differentiation and proliferation^30^. *Lrch4* encodes toll-like receptor (TLR) accessory proteins that regulate the innate immune response^31^. *Serpinb5* plays a role in regulation of macrophage phenotypes^32^. Downregulated *Snx31* encodes integrin α5β1 proteins which mediates cell adhesion to the extracellular matrix by binding fibronectin^33^. Notably, *Fbn1* was upregulated in uPA-CBP mice but did not meet FDR significance level. *Stk36* impacts motile cilia lining on ependymal cells of the nervous system that affect cerebrospinal fluid (CSF) flow^34^. *Rmi2* was upregulated and its expression accelerates the proliferation and migration of cells via activation of the PI3K/AKT pathway in breast cancer models^35^.

### Nociceptive effects of leukocytes in the DRG are activation state- and subtype-specific

Leukocytes are essential components of the immune system and are categorized into granular neutrophils, agranular lymphocytes (B cells, T cells, and natural killer cells) and monocytes which differentiate into macrophages or dendritic cells to aid in phagocytosis and antigen presentation. Our findings suggest increased counts of immune cells that are of myeloid lineage such as monocytes recruited to DRG tissue in uPA-CBP mice compared to sham controls. More established neuropathic pain models have shown increased expression of CD11b+ myeloid-monocyte cells within DRG tissue compared to controls^22,36^. CD11b is a marker commonly expressed on myeloid cells, particularly on monocytes, macrophages, and granulocytes^37^. Our data support that CD45+CD11b+ monocytes were significantly accumulated in uPA-CBP mice compared to sham controls. Neutrophils also express high levels of CD11b but are negative for classical and non-classical macrophage markers such as CD14 and F4/80. Flow cytometric analysis confirmed increased count and percentage of CD45+CD11b+CD14+ and CD45+CD11b+CD14-cell populations, indicating a mixed presence of neutrophils and macrophages. Visual inspection of IHC images determined that CD11b+ cell populations reside near peripherin-labeled neuronal bodies and are largely monocyte-macrophages due to their size and agranularity. Tissue-resident macrophages were also with macrophage surface marker F4/80. Additionally, CD11b co-stained with F4/80-expressing cells which further supports that the cell population is likely a combination of circulating and tissue-resident macrophages. While CD45+CD11b+ populations primarily represent myeloid cells, excluding lymphocytes, dendritic cells, and progenitor cells, CD3, a key component of the T-cell receptor (TCR) complex, is specifically expressed on lymphocytes^38^; however, the CD45+CD11b-CD3+ cell count and its percentage relative to the parent population were negligible, indicating minimal T-cell presence in the DRG tissue.

Interestingly, macrophage accumulation precedes SGC activation^39^ which was implicated in the uPA-CBP DRG by significantly increased intracellular GFAP+TNFα+ cell count and percentage compared to shams. These immune recruitment cascades are dependent on which pathway activation and immune receptor gene expression which has been shown to be subtype-specific and dependent on neuronal subtype as well^40^.

Neurons can direct and influence immune cells directly as well as respond to inflammatory immune agents^21,41,42^. Nociceptors can influence inflammation and immune responses through the release of neuropeptides such as substance P and CGRP^41,43^. For example, Fattori et al 2024 found that in an endometriosis model of chronic pain, nociceptor ablation reduced pain, monocyte recruitment, and lesion size, suggesting that nociceptor activation and neuropeptide release influence the neuronal and non-neuronal environment^43^.

After finding that CD45+CD11b+ cells were highly expressed in uPA-CBP male mice, we performed an experiment to isolate the effects of monocyte-derived cells by administering a CSF1R antagonist, PLX5622, which prevents immature myeloid cells from being recruited at the time of injury by blocking their differentiation into macrophages 3 days prior to uPA injection^18^. The purpose was to study recruited monocyte-derived macrophage populations’ influence on allodynia in mice injected with uPA. DRG-resident macrophage proliferation also depends on CSF1R activation to a lesser degree^39^. For this reason, we chose to use PLX5622 to test if DRG macrophages play a role in maintaining allodynia over time. Neutrophils are unaffected by CSF1R activity. Since the mice showed only reduced mechanical hypersensitivity instead of total recovery, it can be inferred that monocyte-derived macrophages are contributing to mechanical hypersensitivity.

DEGs involved in regulation of the NF-κB pathway were upregulated in uPA-CBP compared to sham controls. For example, *MCCC1* was upregulated and activates the NF-κB pathway which promotes the expression of IL-6^44^. Interestingly, *Alg3* was also significantly upregulated and mediates glycosylation of TGF-β receptor II which is highly implicated in managing inflammation as well as promoting IL-6 in the PPAR-γ/NF-κB pathway^45^. The following DEGs were significantly upregulated and are associated with inflammation. *Ccdc63* encodes proteins linked to moderate association between autoantibodies and muscle/joint pain in a study observing changes post-COVID infection^46^. *Ndor1* encodes electron transfer proteins in the somatosensory system that generate reactive oxidative species (ROS)^47^.

### Limitations

Clinical studies show sex differences in transcriptomic profiles of serum proteins of patients with persistent low back pain^48^. Determining the role of sex as a biological factor in the development of CBP merits future experiments. However, uncovering specific sex-dependent mechanisms was not the goal of this study. Secondly, flow cytometry experiments surveyed CD45+CD11b+ cell populations in DRG of sham and uPA-CBP mice but neuroimmune interactions in DRG have been found to be activation state- and subtype-specific. Additionally, CNS immune and glial cells like microglia and astrocytes similarly contribute to nociceptor sensitization through interactions at the central termination sites of the peripheral afferents^21,49^. However, these CNS cell populations were not tested in this study.

### Conclusions

This study highlights the importance of the DRG neuroimmune axis in chronic back pain. Our uPa-induced chronic back pain model draws attention to the crosstalk between immune cells and sensory neurons that may be maintaining chronic back pain. We also reveal novel targets for the potential development of non-opioid therapeutics for the treatment of chronic back pain.

## Acknowledgments

We thank Dr. Geoffroy Laumet and his graduate student Sim Jaewon at Michigan State University for their valuable feedback on initial flow cytometry experiments.

